# Ribosomal protein mutation suppresses gonadal leader cell migration defects in *mig-17/ADAMTS* mutants in *Caenorhabditis elegans*

**DOI:** 10.1101/2023.03.28.534640

**Authors:** Hon-Song Kim, Kaito Mitsuzumi, Shohei Kondo, Rie Yamaoka, Shinji Ihara, Hiroshi Otsuka, Yukihiko Kubota, Toshinobu Fujiwara, Yukimasa Shibata, Kiyoji Nishiwaki

## Abstract

The migration of the gonadal distal tip cells (DTCs) in *Caenorhabditis elegans* provides an excellent model for studying the migration of epithelial tubes during organogenesis. Mutations in the *mig-17/ADAMTS* gene cause misdirected migration of DTCs during gonad formation, resulting in deformed gonad arms. An amino-acid substitution in RPL- 20 corresponding to the mammalian RPL18a/eL20, a component of the 60S ribosomal large subunit, showed a slow growth phenotype and strongly suppressed the *mig-17* gonadal defects. Slow-growing mutants *clk-1* and *clk-2* also suppressed *mig-17*, although weaker than *rlp-20* mutants. MIG-17 recruits FBL-1C/fibulin-1C to the gonadal basement membrane to regulate DTC migration. Reducing the gene dosage of *fbl-1* by half partially compromised the suppressor activity of the mutant *rpl-20* gene on *mig-17*. Analysis using the mNeonGreen-FBL-1 reporter revealed that its localization to the gonadal basement membrane was significantly reduced in *mig-17*, whereas it was recovered to the wild-type levels in *mig-17; rpl-20* double mutants. These results indicate that the *rpl-20* mutation suppresses *mig-17* gonadal defects through dual mechanisms: deceleration of growth rate and enhancement of FBL-1C recruitment to the gonadal basement membrane.

## Introduction

The ADAMTS (a disintegrin and metalloprotease with thrombospondin motifs) family of secreted zinc metalloproteases plays important roles in morphogenetic processes during animal development. ADAMTS proteases often degrade extracellular matrix proteins in these processes (APTE 2009). For example, ADAMTS-9 and -20 cleave the proteoglycan versican and are required for digit formation and closure of the palate in mice (MCCULLOCH *et al*. 2009; ENOMOTO *et al*. 2010). ADAMTS-1 is also a versicanase, and the *ADAMTS-1* null mice exhibit growth retardation with malformation of adipose, ureteral and adrenal tissues (SHINDO *et al*. 2000; RUSSELL *et al*. 2003; MITTAZ *et al*. 2004). However, the precise roles of ADAMTS proteases in development still remain elusive.

An ADAMTS protease, MIG-17, in *C. elegans*, is involved in the directional regulation of gonadal leader cells, the distal tip cells (DTCs), during the formation of the U-shaped gonad arms in larval development. In the *mig-17* mutants, the misdirected migration of DTCs leads to the deformation of the gonad arms (NISHIWAKI *et al*. 2000). Through genetic suppressor analyses, several gain-of-function amino acid substitutional mutations were identified in extracellular matrix protein genes, including *fbl-1/fibulin-1*, *emb-9/collagen IV a1 chain*, and *let-2/collagen IV a2 chain* (KUBOTA *et al*. 2004; KUBOTA *et al*. 2008; KUBOTA *et al*. 2012; IMANISHI *et al*. 2020). Genetic and molecular analyses of these suppressor mutations revealed the regulatory pathway by which MIG- 17 recruits fibulin-1 to the basement membrane and activates fibulin-1 and collagen IV to modulate downstream events affecting the directional migration of DTCs (KUBOTA *et al*. 2008; IMANISHI *et al*. 2020). Since MIG-17 is a matrix metalloprotease, it is reasonable that genetic suppressors were found in genes encoding extracellular matrix proteins.

In the present study, we identified a novel suppressor mutation, *tk73*, in the ribosomal protein gene *rpl-20*, which corresponds to *RPL18a/eL20*, one of the components of the 60S large ribosomal subunit (BAN *et al*. 2014). The suppressor *rpl- 20(tk73)* mutation was an amino acid substitution and acted as a gain-of-function allele to suppress the *mig-17* DTC migration abnormality. The amount of 60S large ribosomal subunit was markedly reduced in the *rpl-20(tk73)* mutants. Although the decelerated larval growth rate of *rpl-20(tk73)* partly accounted for the mechanism of suppression, it could not fully explain the strong suppression phenotype. MIG-17 regulates DTC migration through sequential recruitment of FBL-1C/fibulin-1C and NID-1/nidogen to the gonadal basement membrane (KUBOTA *et al*. 2008). We found that *mig-17* suppression of *rpl-20(tk73)* depended on extracellular matrix proteins NID-1 and FBL-1. The reduced basement membrane accumulation of FBL-1 in *mig-17* mutants was recovered to wild-type levels in the *rpl-20(tk73)* mutants, suggesting that *rpl-20(tk73)* suppresses the gonadal defect of *mig-17* through promoting basement accumulation of FBL-1C as well as decelerating the larval growth rate. While ribosomal protein mutations are generally thought to affect translation efficiency, defects in ribosome biosynthesis caused by mutations in single ribosomal protein genes lead to a tissue-specific phenotypic abnormality called ribosomopathy in humans (KANG *et al*. 2021). Our findings should help to understand the molecular mechanism underlying ribosomopathies.

## Materials and Methods

### Strains and genetic analysis

Culture, handling, and ethyl methanesulfonate (EMS) mutagenesis of *C. elegans* were conducted as described (Brenner 1974). The following mutations and balancers were used in this work: *mig-17(k135, k169, k174), mig-18(k140, gm321), fbl-1(tk45), nid- 1(cg119), unc-42(e270), unc-13(e1091), unc-32(e189), unc-119(e2498), clk-1(gm30)*, *clk-2(qm37)* and *hT2[qIs48] (I; III), nT1[qIs51] (IV; V)*(BRENNER 1974; MADURO AND PILGRIM 1995; WONG *et al*. 1995; NISHIWAKI 1999; BENARD *et al*. 2001). *atf-4(tm4397)* was obtained from National Bioresource Project for the nematode.

### Microscopy

Gonad migration phenotypes were scored using a Nomarski microscope (Axioplan 2; Zeiss). Analysis of gonadal phenotypes was performed at the young-adult stage as described (NISHIWAKI 1999). Confocal laser scanning microscopy was conducted with LSM5 (Zeiss) controlled by PASCAL version 3.2 SP2 or ZEN software (Zeiss) to capture mCherry images. The fluorescence intensities of mNeonGreen were quantified as follows. For each sample, confocal images of a sagittal section of the pharynx were obtained with a Zeiss Imager M2 microscope equipped with a spinning-disk confocal scan head (CSU-X1; Yokogawa) and an ImageEM CCD camera (ImageEM; Hamamatsu Photonics). Using MetaMorph software, the average fluorescence intensities of the boxed areas shown in Figure 8A were measured. The maximum background intensities inside the pharynx were subtracted from each values.

### Molecular identification of rpl-20(tk73)

The *tk73* mutation was isolated as a genetic suppressor of the gonadal defect of *mig-17(k174)* mutants using EMS mutagenesis (BRENNER 1974). Single-nucleotide polymorphism mapping experiments (WICKS *et al*. 2001) placed *tk73* between a 4581 ∽4788kb region on linkage group IV. Whole-genome sequence analyses comparing *mig-17(k174)* and *tk73; mig-17(k174)* genomes identified a single mutation in the coding region of the *rpl-20* gene (<u>G</u>GA to <u>A</u>GA), resulting in the amino acid substitution G82R. Microinjection experiments using a PCR-amplified mutant *rpl-20* gene fragment successfully rescued the gonadal defects of *mig-17* mutants, confirming that *rpl-20* is the gene responsible for *tk73*.

### Constructs

To produce the *rpl-20p::rpl-20(tk73)* plasmid, the genomic region containing 1190 bp 5’-untranslated region (UTR) to 591 bp 3’-UTR of the *rpl-20* gene was PCR amplified from genomic DNA of *mig-17(k174); rpl-20(tk73)* and cloned the fragment into the *Not*I and *Acc65*I sites of pBluescript II KS(-). To produce the plasmids for tissue-specific expression, the 5’-UTR region was replaced with PCR amplified 5’-UTR fragments of *mig-24* (1156 bp), *myo-2* (1463 bp), *myo-3* (2452 bp) or *elt-2* (5044 bp). *mig-24p::rpl-20(WT) and let-2p::rpl-20(WT)* plasmid were generated by site-directed mutagenesis of *mig-24p::rpl-20(tk73) and let-2p::rpl-20(tk73)* plasmids, respectively.

The *rpl-20p::mCherry::rpl-20(WT)* plasmid was constructed using pPD95.79 in which the coding region of *GFP* was replaced with that of *mCherry*. *fbl-1C cDNA::Venus* plasmid was constructed form the *fbl-1C::3HA(k201, ΔD)* plasmid (KUBOTA *et al*. 2004) by replacing the fragment from the fourth exon to 3HA into *fbl-1C* cDNA.

### Germline transformation

Germline transformation was carried out as described (Mello *et al*. 1991). Transgenic strains were made by injecting plasmids into *unc-119(e2498)* hermaphrodites and the generated transgenic arrays were transferred to appropriate genetic backgrounds having *unc-119(e2498)* by mating. *mig-24p::rpl-20(tk73), myo-3p::rpl-20(tk73), elt- 2p::rpl-20(tk73), mig-24p::rpl-20(WT), myo-3p::rpl-20(WT),* and *elt-2p::rpl-20(WT)* plasmids were injected at 10 ng/μl with 30 ng/μl *unc-119^+^*plasmid (pDP#MM016B)

(MADURO AND PILGRIM 1995) and 110 ng/μl pBluescriptII KS(–) (carrier DNA). *myo-2p::rpl-20(tk73)* plasmid was injected at 2 ng/μl with 38 ng/μl *unc-119^+^* plasmid (pDP#MM016B) and 110 ng/μl pBluescriptII KS(–).

### Western blot analysis

Western blot analysis was done as described (IHARA AND NISHIWAKI 2007) using anti-NID-1 (2 μg/ml) (KUBOTA *et al*. 2008) and anti-α-tubulin (DM1A, 1:1000; abcam), anti-GFP (3E6, 1:200; Molecular Probes) or anti-mCherry (5f8 1:1000, Funakoshi).

### Ribosome assay

Preparation of worm samples and sucrose density gradient centrifugation were conducted as described (SAIJOU *et al*. 2004). Sucrose cushion centrifugation was conducted as described (FUJIWARA *et al*. 2001).

## Results

### An amino acid substitution in the ribosomal protein gene *rpl-20* suppresses the abnormal distal tip cell migration of *mig-17* mutants

In *C. elegans*, the distal tip cells (DTCs) located at the anterior and posterior ends of the gonad primordium act as leader cells to elongate and form the U-shaped gonad arms. The ADAMTS family metalloprotease MIG-17 is required for the directional migration of DTCs (NISHIWAKI *et al*. 2000). The *mig-17* mutants show deformed gonad formation due to meandering and straying of DTCs (NISHIWAKI 1999). Through forward genetics screening, we identified a mutation *rpl-20(tk73)* that could alleviate the gonadal defect of *mig-17* mutants (Figure 1, A and B). RPL-20 was an ortholog of the mammalian ribosomal protein RPL18a/eL20, a component of the 60S large ribosomal subunit (BAN *et al*. 2014). *rpl-20(tk73)* was an amino acid substitution of evolutionarily conserved glycine 82 (glycine 79 in mammals) into arginine (Figure 1C). It suppressed the *mig- 17(tk174)* null allele strongly as a homozygote and weakly as a heterozygote, while alone showed no gonadal defects. The deletion allele *rpl-20(ok2256)*, which totally deletes the coding region, exhibited a homozygous early larval lethal phenotype and *rpl- 20(ok2256)/+* failed to suppress *mig-17(tk174)*, implying that *rpl-20(tk73)* is a gain-of- function allele (Figure 1B). *rpl-20(tk73)* also suppressed missense *mig-17* alleles *k135* and *k196*, as well as mutations in *mig-18* that encodes a cofactor for the MIG-17 metalloprotease (KIM *et al*. 2014), although the suppression for *mig-18* appeared to be weaker than for *mig-17* (Figure 2A). However, it failed to suppress the gonadal defects caused by RNAi knockdown of *gon-1*, another ADAMTS metalloprotease that functions in gonadogenesis (HESSELSON et al. 2004). Similarly, *rpl-20(tk73)* could not suppress the *mig-17*-like meandering DTC migration defects caused by mutations in *sqv-5* (chondroitin synthase) and *mig-22* (chondroitin polymerizing factor) (SUZUKI *et al*. 2006) (Figure 2, B and C). These results suggest that *rpl-20(tk73)* specifically suppresses the gonadal defects caused by mutations in *mig-17* and its cofactor *mig-18*.

**Figure 1.**
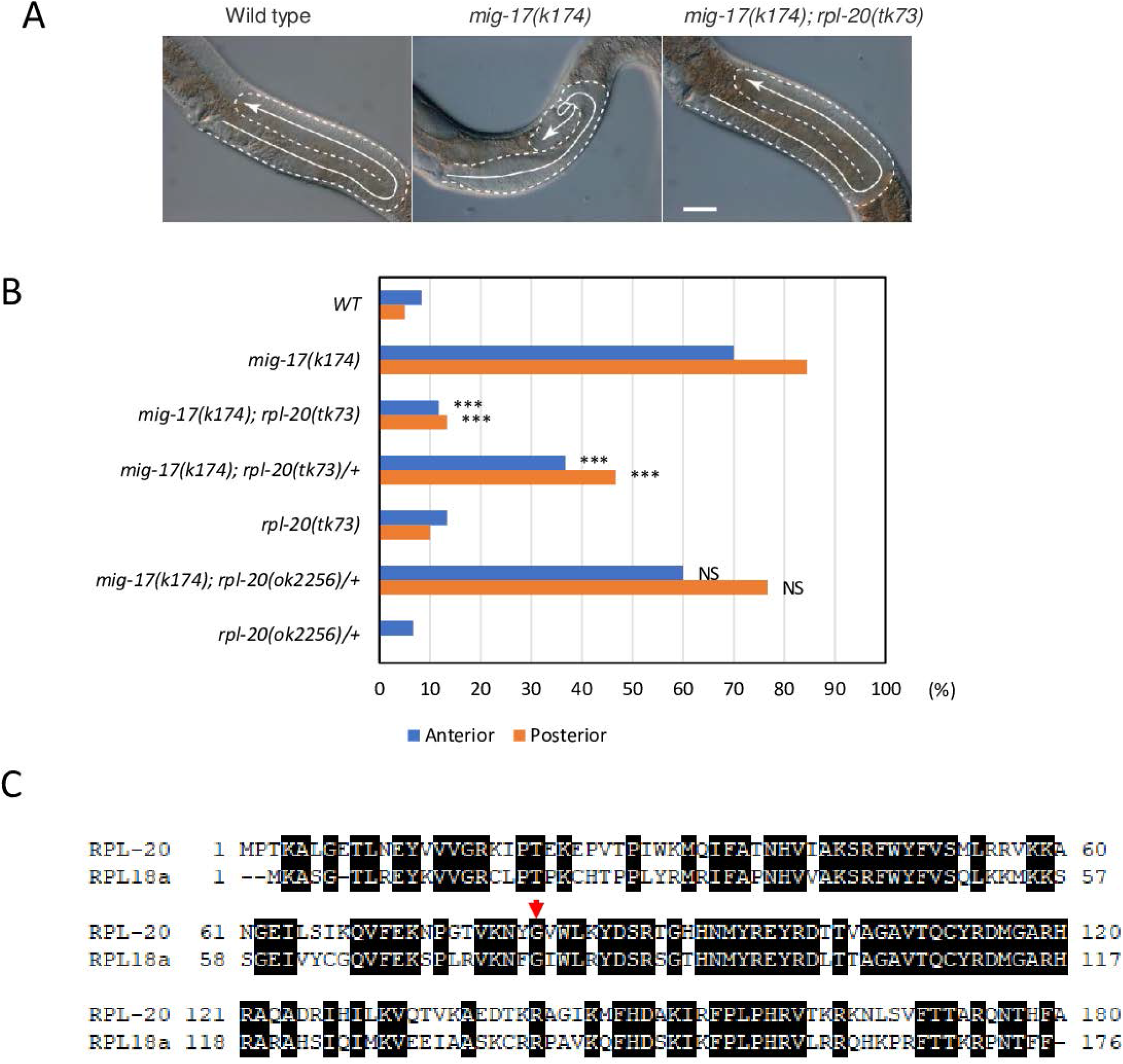
Suppression of *mig-17* DTC migration defects by *rlp-20(tk73)*. (A) Gonad morphology (arrows) of wild type, *mig-17(k174),* and *mig-17(k174); rpl-20(tk73)* young-adult hermaphrodites. Posterior gonads are shown. Anterior to the left, dorsal to the top. Bar, 20 μm. (B) Quantitative analysis of gonadal defects. N=60 for each strain. P-values for Fisher’s exact test against *mig-17(k174)* for *mig-17(k174)* carrying strains: ***, P < 0.001; NS, not significant. (C) Amino acid sequence homology between RPL-20 and human RPL81a. Identical amino acids are shown in black boxes. G82 mutated in rpl-20 is depicted with an arrow.

**Figure 2.**
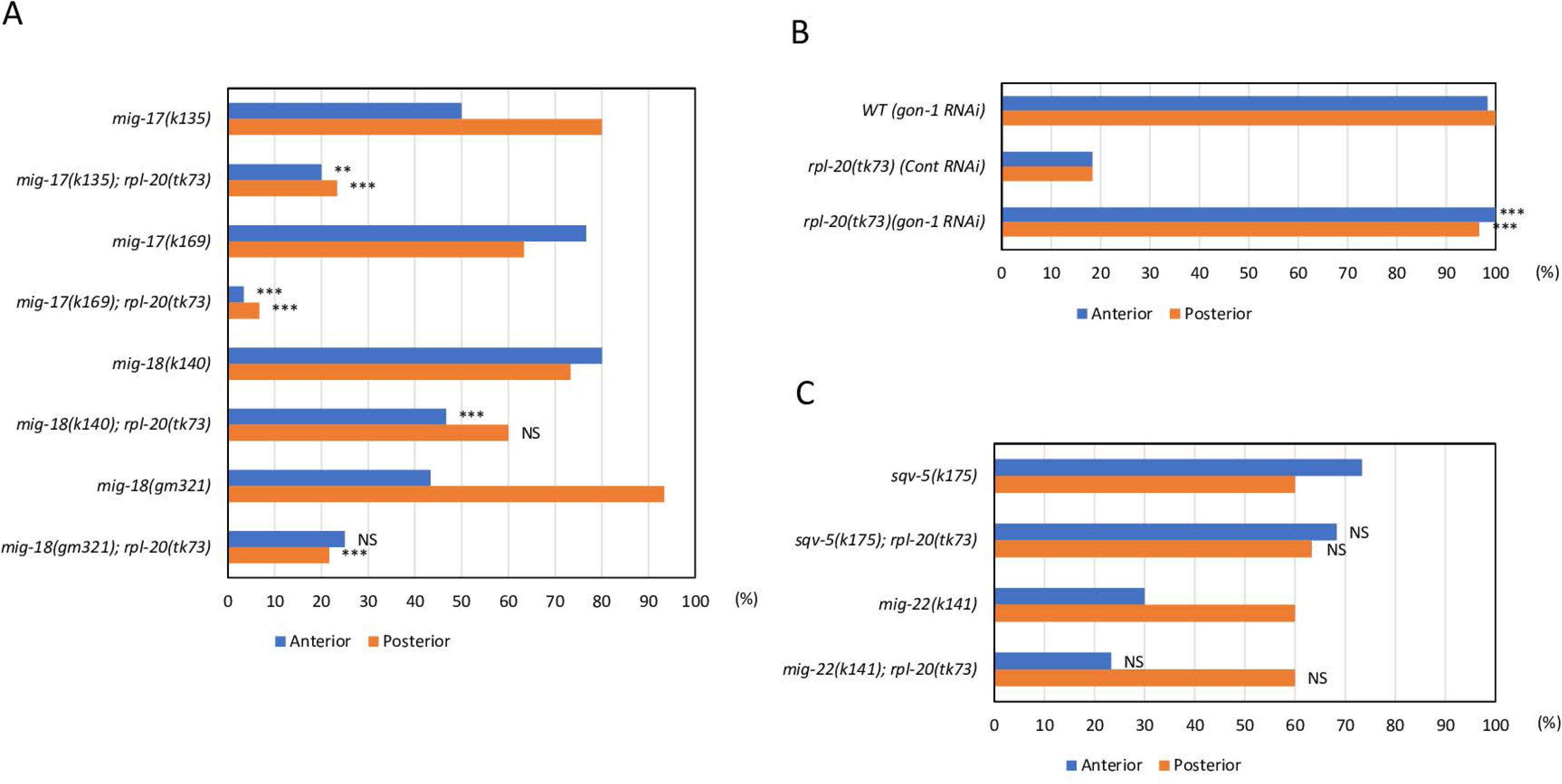
Suppression of *mig-17* and *mig-18* mutants by *rpl-20(tk73)*. (A-C) *rpl-20(tk73)* suppresses the DTC migration defects of *mig-17* and *mig-18* alleles (A), but not *gon-1(RNAi)* (B) or *sqv-5(k175)* and *mig-22(k141)* mutants (C). *sqv-5(k175)* and *mig-22(k141)* were marked with *unc-13(e1091)* and *unc-32(e189)*, respectively. N=60 for each strain. P-values for Fisher’s exact test against *mig-17, mig-18, rpl-20(tk73)* control RNAi*, sqv-5,* and *mig-22* for the respective double mutants: ***, P < 0.001; **, P< 0.01; NS, not significant.

It is reported that the guidance molecule semaphorin regulates epidermal morphogenesis by stimulating general translation through repression of the phosphorylation of eukaryotic initiation factor 2α (eIF2α) and inhibition of the target of rapamycin complex 2 (TORC2) pathway (NUKAZUKA *et al*. 2008; NUKAZUKA *et al*. 2011). We suspected that MIG-17 may also affect translation to regulate DTC migration. This possibility seemed consistent with the fact that a mutation in a ribosomal protein suppresses the *mig-17* defects. Semaphorin signaling promotes eIF2α dephosphorylation by repressing the activities of GCN-1 and PEK-1 (NUKAZUKA *et al*. 2008). Semaphorin also represses RICT-1 (TORC2 component) (NUKAZUKA *et al*. 2011). We generated *gcn-1; mig-17*, *pek-1; mig-17* and *rict-1; mig-17* double mutants and found that they failed to suppress the *mig-17* gonadal defects (Figure S1), suggesting that MIG-17 does not control DTC migration through the stimulation of general translation.

### Growth rate deceleration suppresses the gonadal abnormality of *mig-17* mutants

During the analysis of the *rpl-20(tk73)* mutants, we observed that they grew slower compared to the wild type or *mig-17* mutant animals. *rpl-20(tk73); mig-17(k174)* double mutants also grew slowly. We analyzed their growth rates based on the stages of vulval development (Figure 3, A and B). To investigate whether the growth retardation caused *mig-17* suppression, we created double mutants between *mig-17* or *mig-18* alleles and slow-growing mutants *clk-1(gm30)* or *clk-2(qm37)* (WONG *et al*. 1995; LAKOWSKI AND HEKIMI 1996). We found that these *clk* mutants partially suppressed *mig-17* and *mig-18* gonadal defects (Figure 4, A and B). *clk-1* encodes a mitochondrial hydroxylase that is necessary for ubiquinone biosynthesis, while *clk-2* encodes a component of DNA damage response and telomere metabolism (AHMED *et al*. 2001; BENARD *et al*. 2001), as well as the nonsense-mediated mRNA decay pathway (GUO *et al*. 2021). Thus, these genes have no functional relevance to *rpl-20*, indicating that the slow growing phenotype of these mutants is at least partly contribute to the suppression. Since *clk- 2(qm37)* exhibited a growth delay similar to *rpl-20(tk73)* and *clk-1(gm30)* animals grew much slower than *rpl-20(tk73)* (Figure 3, A-C), it is possible that a mechanism other than growth retardation is responsible for *mig-17* suppression by *rpl-20(tk73)*.

**Figure 3.**
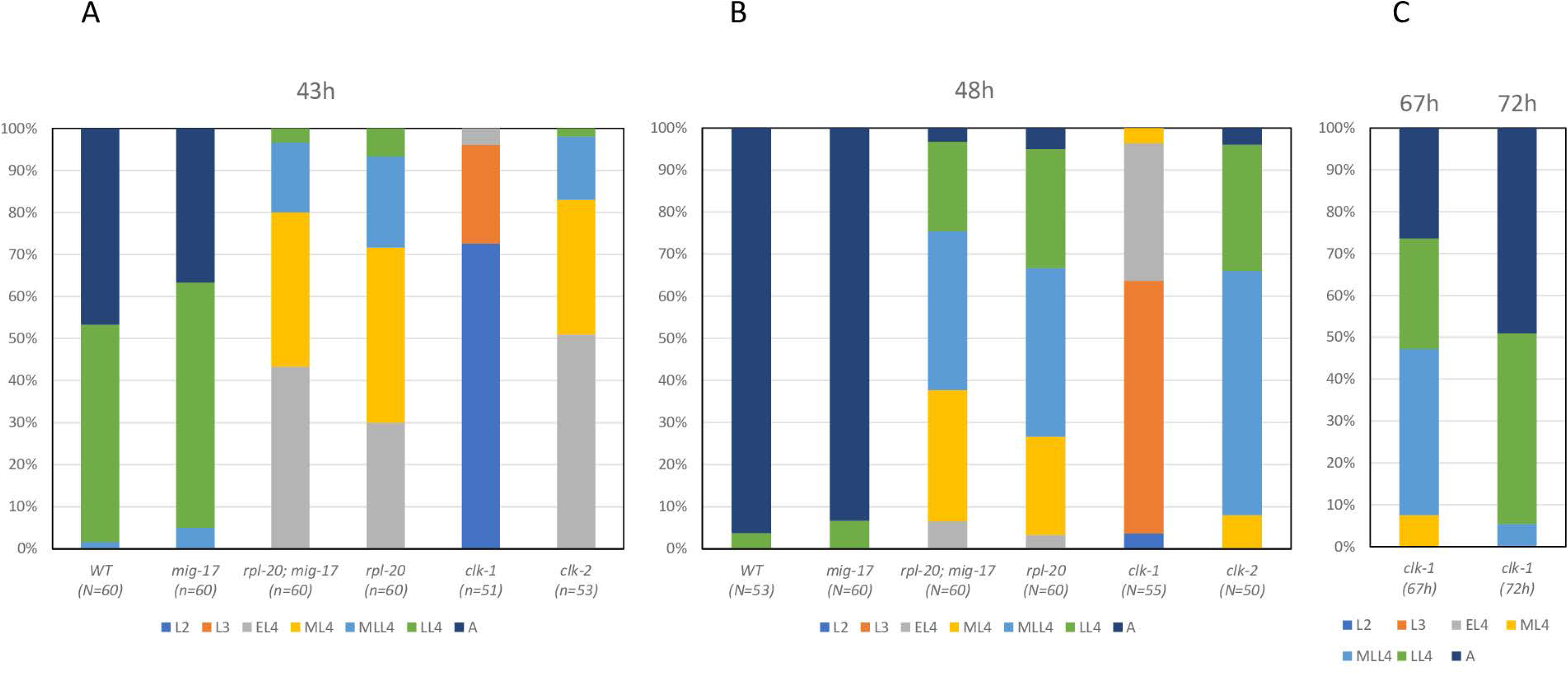
Growth rate analysis of *mig-17, rpl-20, mig-17; rpl-20, clk-1, clk-2* mutants. Larval stages were determined as described by MacNeil et al. (2013). Color codes represent larval stage 2 (L2), larval stage 3 (L3), early L4 (EL4), mid-L4 (ML4), late L4 (LL4), and adult (A), respectively.

**Figure 4.**
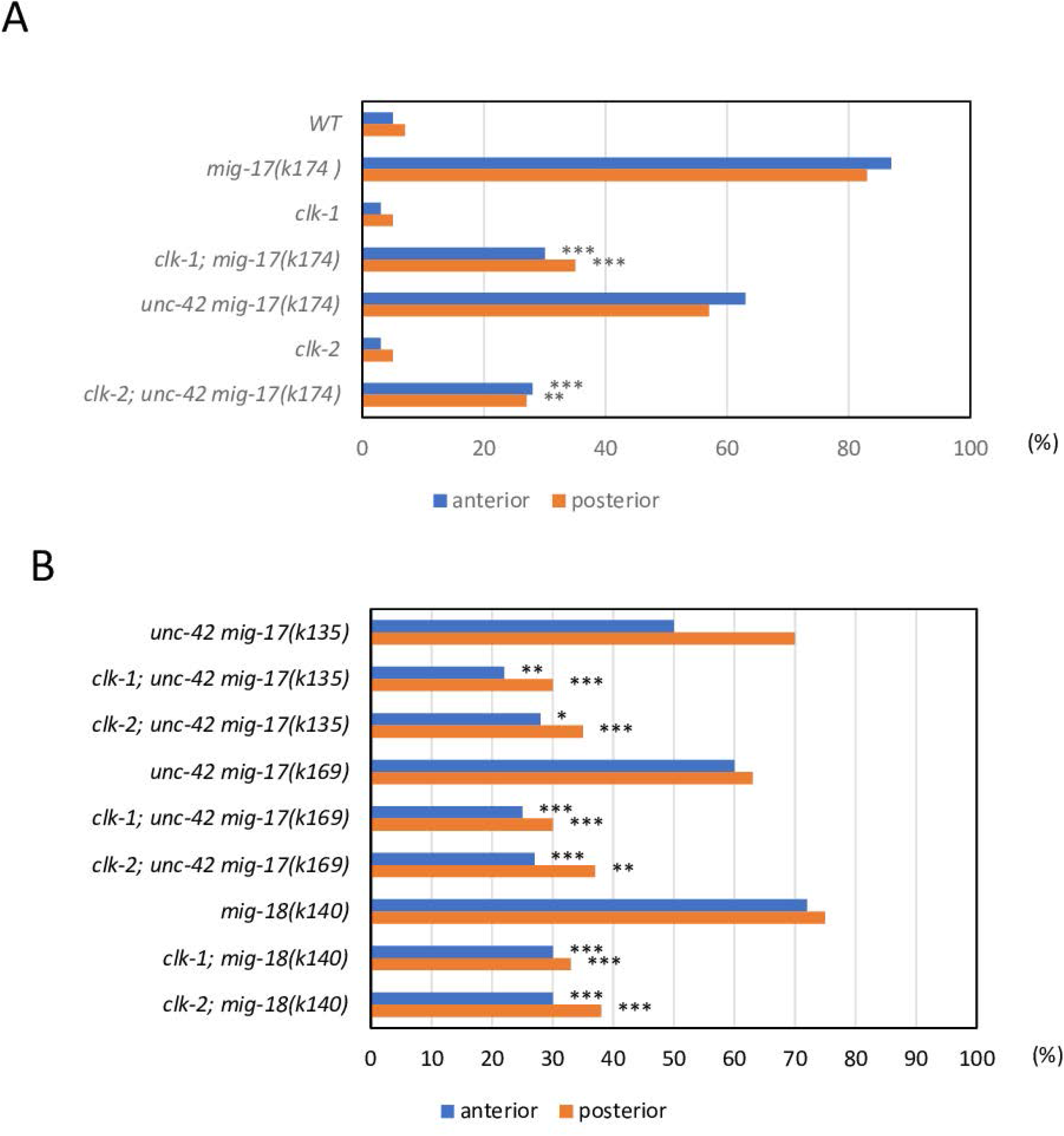
Suppression of *mig-17* and *mig-18* mutants by *clk-1* and *clk-2*. (A) Suppression of *mig- 17(k174)*. (B) Suppression of *mig-17* and *mig-18* alleles. N=60 for each strain. P-values for Fisher’s exact test against *mig-17* and *unc-42 mig-17* for the respective double mutants: ***, P < 0.001; **, P< 0.01; *, P<0.05.

### Intestine-specific expression of mutant RPL-20 efficiently suppresses *mig-17*

We generated a translational fusion construct, *rpl-20p::mCherry::rpl-20(WT)*. This construct was partially functional as it rescued the early larval lethal phenotype of the *rpl-20(ok2256)* deletion allele when introduced as an extrachromosomal array containing multicopy *rpl-20p::mCherry::rpl-20(WT)* transgenes, although the adult animals with the array were sterile. The mCherry expression was detected ubiquitously except in the germline, likely due to gene silencing of the multicopy array (ROBERT *et al*. 2005). The expression appeared most intense in the intestine. The same expression pattern was observed in *rpl-20p::mCherry::rpl-20(tk73)* (Figure 5A).

**Figure 5.**
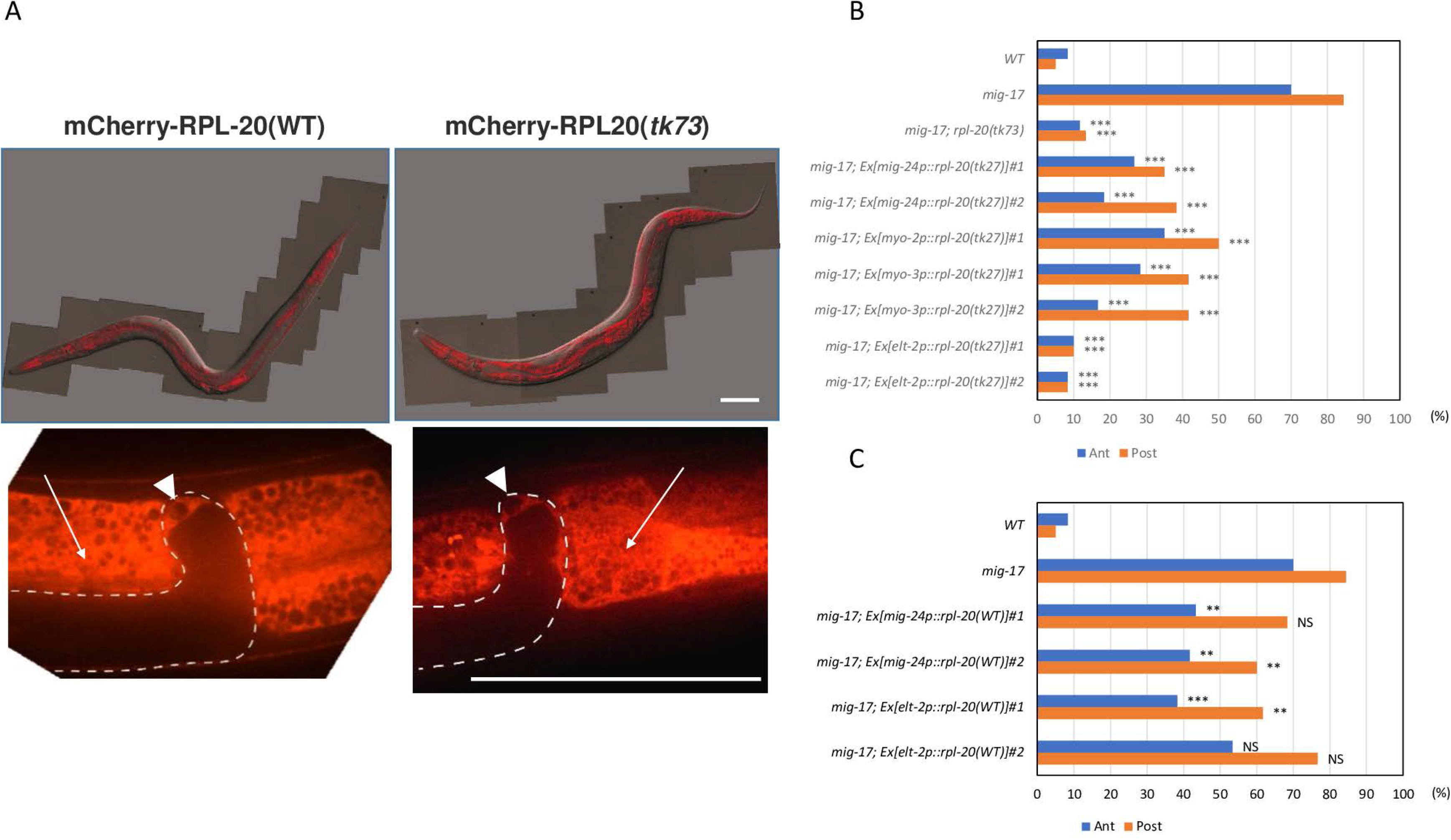
Tissue-specific expression of RPL-20. (A) Expression of mCherry-RPL-20(WT) and mCherry-RPL-20(*tk73*). Upper panels are Nomarski and fluorescence merged images for young adult animals and lower panels represent close-up images of the gonadal tip regions. Arrows and arrowheads indicate intestine and the distal tip cells, respectively. Bars, 50 μm. (B, C) Rescue experiments of the *mig-17* gonadal defect by expressing *rpl-20(tk73)* (B) or *rpl-20(WT)* (C) genes under various tissue-specific promoters. N=60 for each strain. P-values for Fisher’s exact test against *mig-17(k174)* for *mig-17(k174)* carrying strains: ***, P < 0.001; **, P < 0.01; NS, not significant. #1 and #2 are independently established transgenic lines.

To determine the tissues in which expression of the mutant RPL-20(*tk73*) protein is important for suppressing *mig-17*, we expressed the mutant *rpl-20(tk73)* gene under tissue-specific promoters. We found that DTC-specific expression (*mig-24p::rpl- 20(tk73)*) (TAMAI AND NISHIWAKI 2007), pharyngeal muscle-specific expression (*myo- 2::rpl-20(tk73)*) (OKKEMA *et al*. 1993), and body wall muscle-specific expression (*myo- 3::rpl-20(tk73)*) (OKKEMA *et al*. 1993) weakly suppressed *mig-17*, whereas intestine-specific expression (*elt-2p::rpl-20(tk73)*) (FUKUSHIGE *et al*. 1998) strongly suppressed the gonadal defects of *mig-17* mutants (Figure 5B). The suppressor activity of *elt- 2p::rpl-20(WT)* was much weaker than that of *elt-2p::rpl-20(tk73)* (Figure 5C). Next, we analyzed these transgenic lines for their growth rates. Although all these lines exhibited growth retardation, *elt-2p::rpl-20(tk73)* was the slowest, and its growth rate was equivalent to that of the *rpl-20(tk73)* mutant. The growth rate for *elt-2p::rpl- 20(WT)* was slightly slower than that of the wild-type animals (Figure 6, A and B).

**Figure 6.**
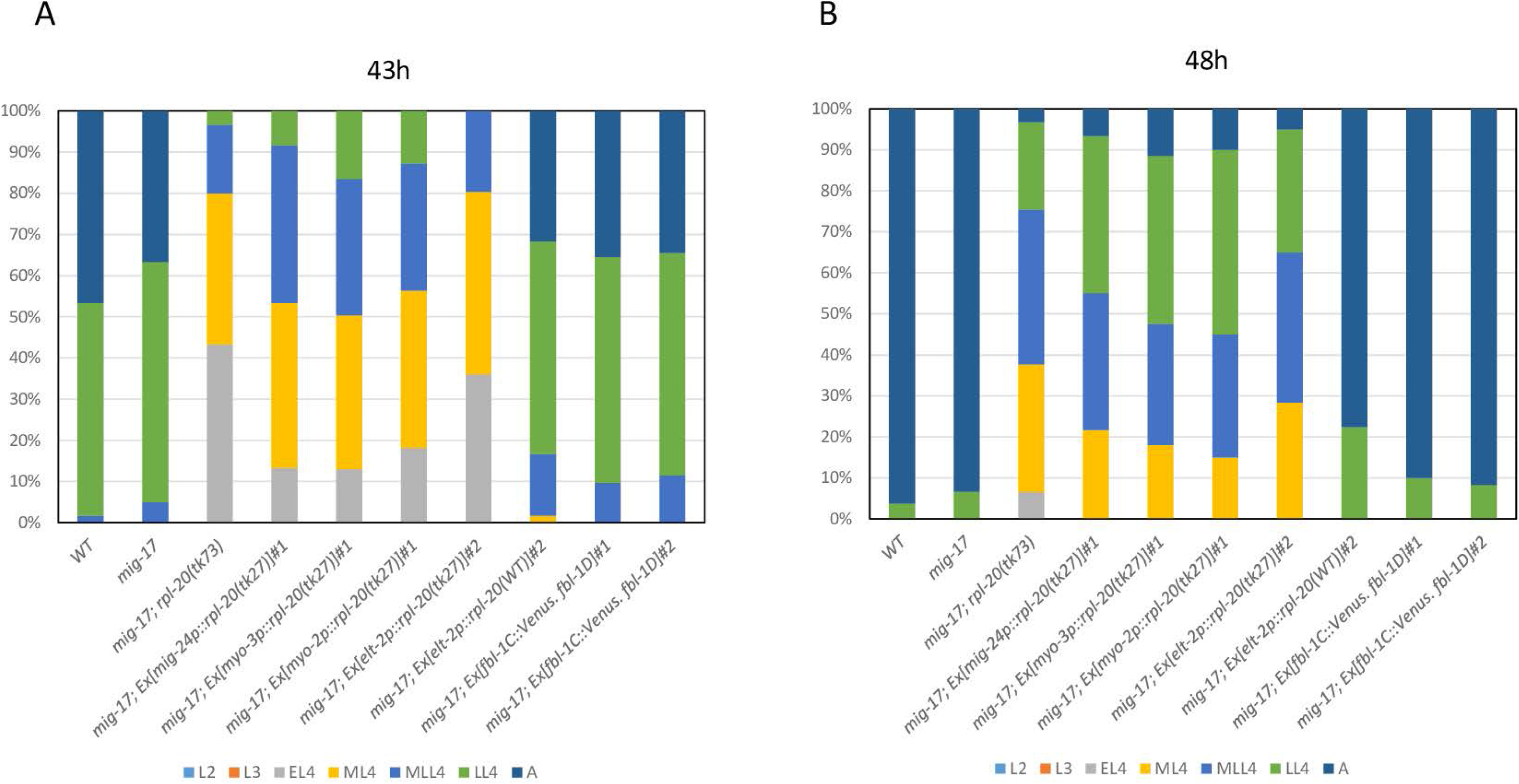
Growth rate analysis of *mig-17* animals carrying various plasmid constructs. Analysis was conducted as in Figure 3. #1 and #2 are independently established transgenic lines.

These results support the idea that the growth rate retardation caused by the *rpl-20(tk73)* allele is causative for *mig-17* suppression and that RPL-20(G82R) expressed in the intestine plays an important role in the suppression.

### mig-17 suppression by rpl-20(tk73) depends on fbl-1 and nid-1

We previously demonstrated that MIG-17 activity is necessary for efficient recruitment and accumulation of FBL-1C to the basement membrane, which in turn recruits NID-1, and that this activity is essential for the directional migration of gonadal DTCs (KUBOTA *et al*. 2008). FBL-1C is a basement membrane protein that is expressed and secreted from the intestine (KUBOTA *et al*. 2004). As *rpl-20(tk73)* expression in the intestine had the strongest effect on *mig-17* suppression, it is possible that FBL-1C and NID-1 are involved in *rpl-20(tk73)* suppression of *mig-17*. To explore this possibility, we introduced a *nid-1(cg119)* null allele into *mig-17(k174); rpl-20(tk73)* double mutants and found that the suppressor activity of *rpl-20(tk73)* was mostly abolished (Figure 7A). We also reduced the *fbl-1* gene dosage by half in the *mig-17(k174); rpl-20(tk73)* mutant background and observed that the suppression was partially compromised. Moreover, overexpression of FBL-1C and D or FBL-1C alone in *mig-17(k174)* animals significantly suppressed their gonadal abnormality (Figure 7B). We observed no difference in the growth rate of these transgenic animals overexpressing *fbl-1* transgenes (Figure 6, A and B). These results suggest that *rpl-20(tk73)* may affect the amount of FBL-1 expression to exert its activity in suppressing *mig-17*, possibly through its mutationally modified translational activity. We compared the levels of NID-1 expression in wild type, *mig-17(k174); rpl-20(tk73)*, and *rpl-20(tk73)* animals using anti-NID-1 immunoblotting and found that the expression levels of NID-1A/B and NID- 1C were similar among these strains (Figure 7C). We also compared the levels of FBL- 1C-Venus expression between wild type and *rpl-20(tk73)* backgrounds using anti-GFP immunoblotting and observed that the amount of FBL-1C-Venus protein was slightly lower in *rpl-20(tk73)* compared to wild type, indicating that the suppressor activity of *rpl-20(tk73)* does not result from the upregulation of NID-1 or FBL-1C expression (Figure 7D).

**Figure 7.**
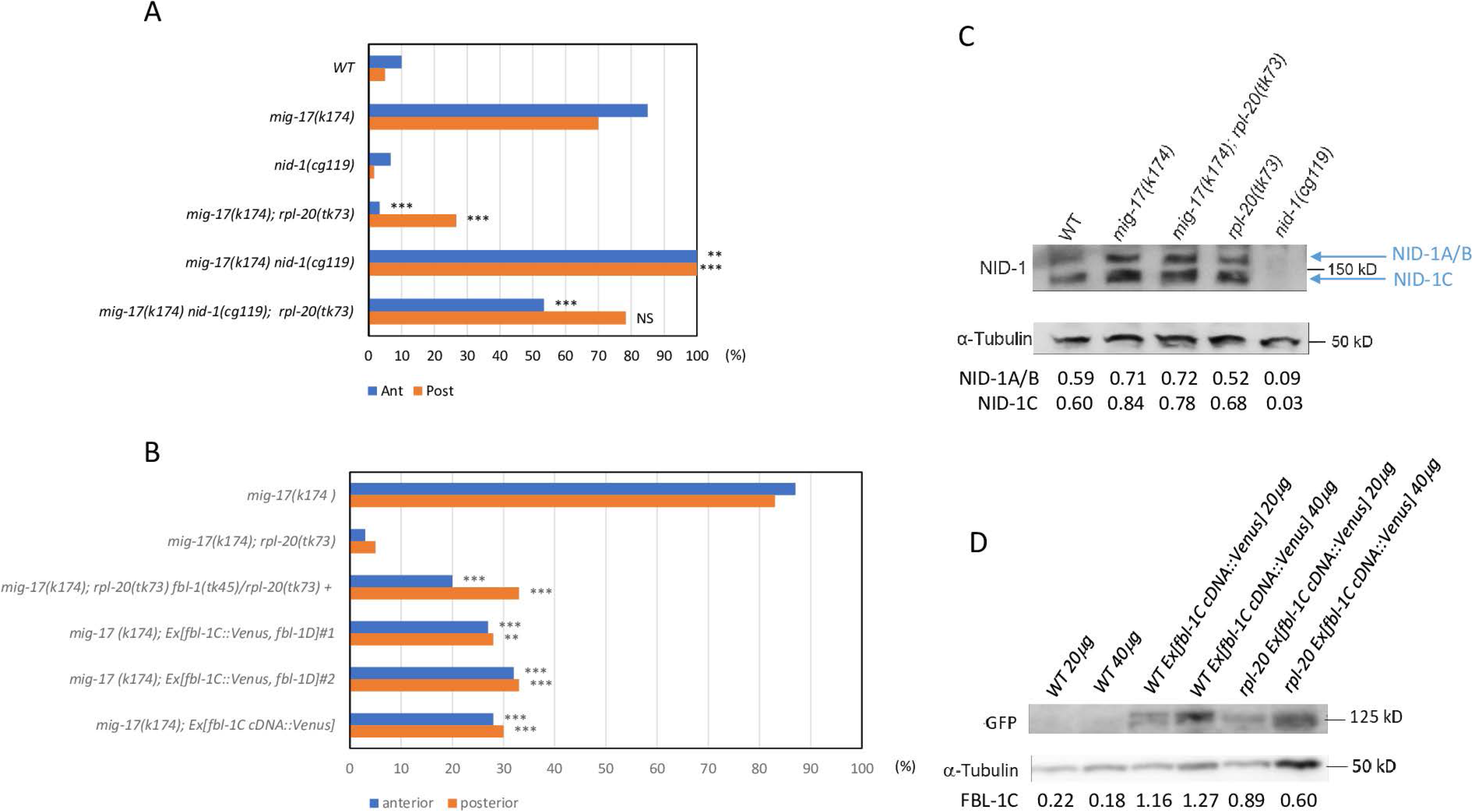
Effects of *nid-1* and *fbl-1* mutations or overexpression on *mig-17* suppression. (A) Effect of *nid-1* mutation. (B) Effect of *fbl-1* mutation or overexpression. *rpl-20(tk73) fbl-1(tk45)* chromosome is marked with *dpy-13(e184)*. N=60 for each strain. P-values for Fisher’s exact test against *mig-17(k174)* for *mig-17(k174)* carrying strains: ***, P < 0.001; **, P < 0.01; NS, not significant. (C) Western blot analysis of NID-1 expression using anti-NID-1 (Kubota et al.,2008). The NID-1A/B (upper), NID-1C (lower) and anti-α-tubulin (loading control) protein levels on western blots were measured by Image J software. The intensity ratios (NID-1/tubulin) were indicated below respective lanes. (D) Western blot analysis of fbl-1C cDNA::Venus using anti- GFP. The FBL-1C-Venus and tubulin protein levels were analyzed as in (c).

### The amount of FBL-1 basement membrane localization is affected in *rpl-20(tk73)*

Using a transgenic line in which the mNeonGreen reporter is placed at the N-terminus of *fbl-1* coding region by the CRISPR/Cas9 method (KEELEY *et al*. 2020) we examined the extracellular matrix (ECM) localization of the FBL-1C and FBL-1D isoforms. The *fbl-1* gene produces various splicing isoforms having the same N-terminus and either one of the two types of C-terminal domains, generating two groups of splicing variants classified as FBL-1C or FBL-1D (KUBOTA *et al*. 2004; MURIEL *et al*. 2005). Although it is known that FBL-1C, but not FBL-1D, localizes to the gonadal basement membrane (KUBOTA *et al*. 2004), the mNeonGreen fluorescence was not detectable in this location, probably because of its accumulation below the detection level. Thus, we examined the levels of mNeonGreen-FBL-1 accumulation in the pharynx. In the pharynx, FBL-1D accumulates along the flexible tracks at the anterior tip of the pharynx, whereas FBL-1C does so in the pharyngeal basement membrane (MURIEL *et al*. 2005). We quantified mNeonGreen-FBL-1 fluorescence intensities using confocal images. We found that the amounts of FBL-1D at the flexible tracks, as well as FBL-1C in the basement membrane, were significantly greater in *mig-17(k174); rpl-20(tk73)* compared to that in *mig-17(k174)* (Figure 8, A-D). These results suggest that *rpl-20(tk73)* affects the localization of FBL-1 proteins to the basement membrane rather than their translational levels. Presumably, FBL-1C accumulation in the gonadal basement membrane, in addition to the pharyngeal basement membrane, should be enhanced in *rpl-20(tk73)* to contribute to the suppression.

**Figure 8.**
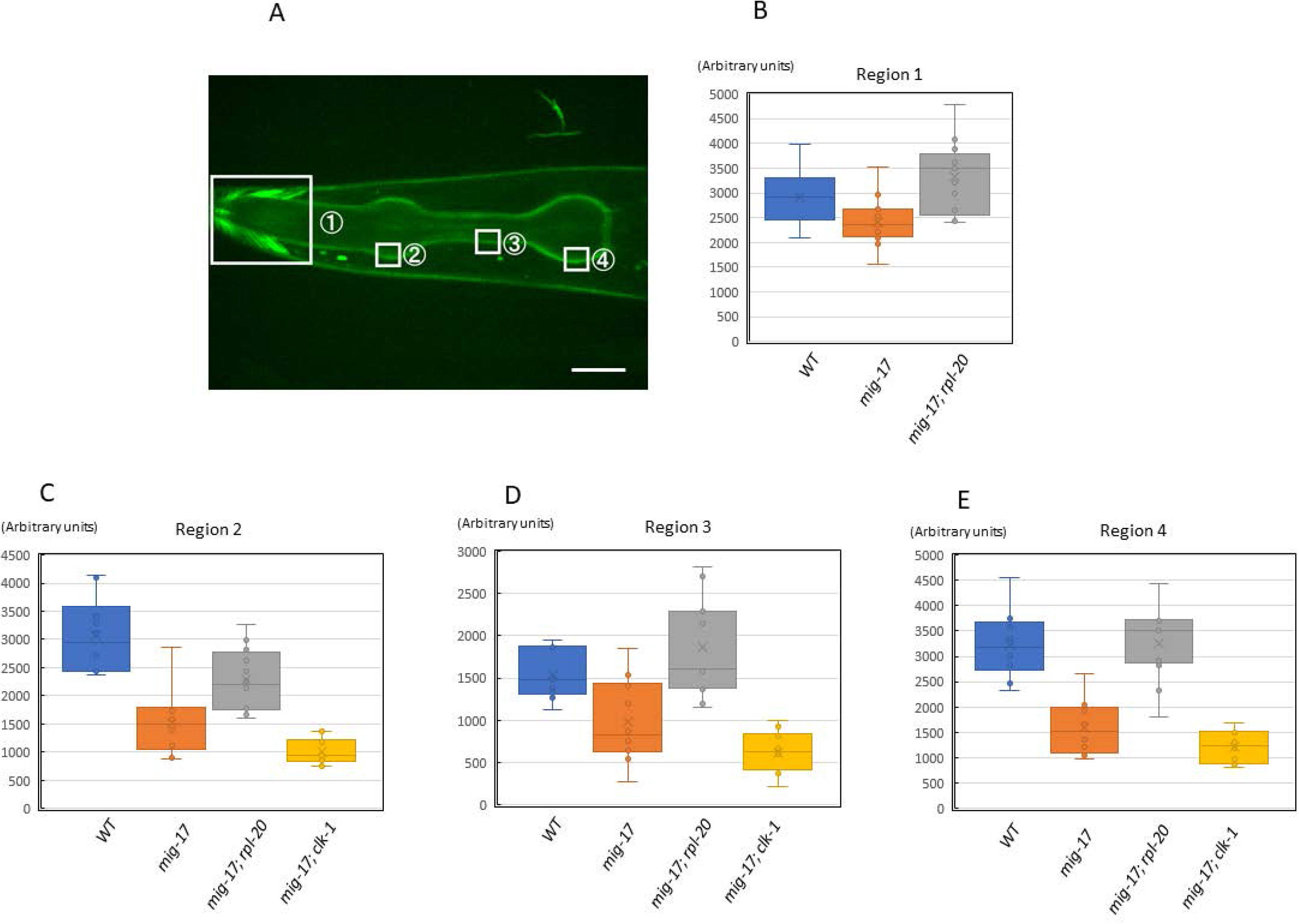
Expression of mNG-FBL-1. (A) Confocal image of the head region of a wild type young adult hermaphrodite expressing mNG-FBL-1. Bar, 20 μm. (B-E) Average fluorescence intensities of the boxed regions were plotted (see experimental procedures). Same sized boxes were used for each experiment. N=11-13.

We then introduced *mNeonGreen::fbl-1* into *mig-17(k174); clk-1(gm30)* mutant background. We observed that the levels of mNeonGreen-FBL-1 accumulation in *clk- 1(gm30)* mutants were not significantly different from that in the *mig-17* mutants (Figure 8, C-E). Taken together, these results suggest that *rpl-20(tk73)* suppresses *mig-17* by decelerating the growth rate as well as enhancing FBL-1C accumulation to the basement membrane.

### Reduction of 60S ribosomal subunit in *rpl-20(tk73)* mutants

Western blot analysis was performed on ribosomes extracted from animals carrying *rpl- 20p::mCherry::rpl-20(WT)* and *rpl-20p::mCherry::rpl-20(tk73)* arrays using anti-mCherry. The results showed that both the wild-type and mutant RPL-20 proteins were incorporated into the ribosome (Figure 9A). We analyzed ribosome profiles by sucrose density gradient centrifugation using extracts from wild type and *rpl-20(tk73)* animals. It was observed that there was a concomitant reduction of the 60S subunit and the 80S ribosomes in *rpl-20(tk73)*, indicating that the RPL-20(*tk73*) mutant protein partially interferes with the biogenesis of the 60S subunit, resulting in a decrease in the amount of 80S mature ribosomes (Figure 9B). It is reported that reduced biogenesis of the 60S subunit can extend yeast lifespan by translational upregulation of the nutrient-responsive transcription factor Gcn4. Therefore, we examined whether the depletion of the *C. elegans* Gcn4 ortholog ATF-4 could abrogate *rpl-20(tk73)*-dependent suppression of *mig-17*. However, it was observed that the deletion mutation *atf-4(tm4398)* had no influence on the suppressor activity of r*pl-20(tk73)* (Figure S2). Thus, it is likely that molecules other than ATF-4 function to transduce the *rlp-20(tk73)* activity to suppress the *mig-17*-dependent gonadal defects.

**Figure 9.**
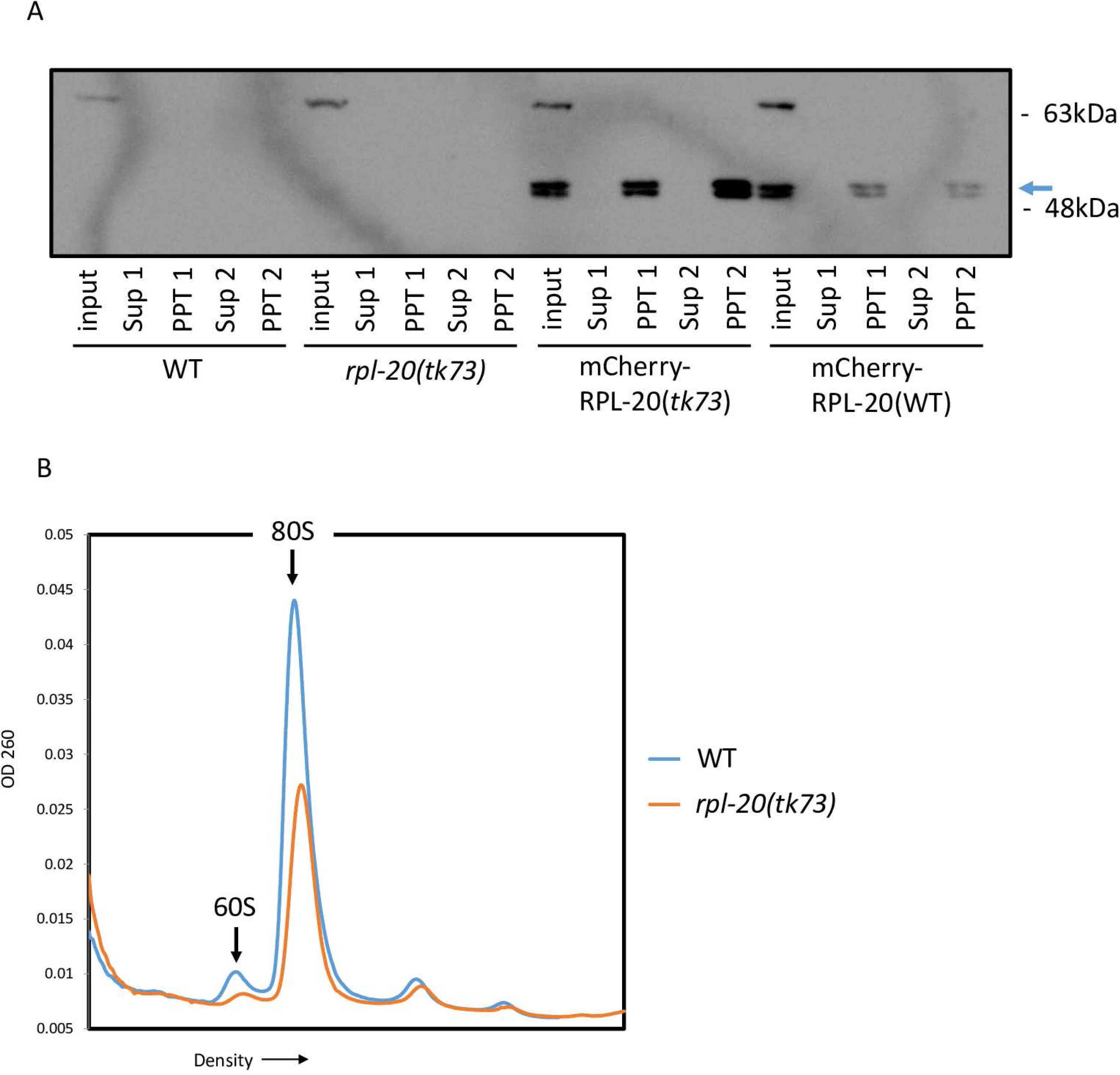
Analysis of ribosomes. (A) Western blot analysis of ribosomes for wild type animals expressing mCherry-RPL-20(*tk73*) or mCherry-RPL-20(WT). Ribosomes were precipitated by dual sucrose cushion centrifugation and the samples of supernatants and precipitates were analyzed. Sup 1 and 2 and PPT 1 and 2 represent supernatant and precipitate samples from first and second centrifugation, respectively. Arrow indicates the bands corresponding to mCherry-RPL-20. The double bands may be generated due to post-translational modification of RPL-20 (SIMSEK AND BARNA 2017). (B) Polysome profiles of wild type and *rpl-20(tk73)* animals. The amounts of 60S subunit and 80S ribosome in *rpl-20(tk73)* were reduced relative to the wild type.

## Discussion

In this study, we identified a novel missense mutation, *rpl-20(tk73)*, that strongly suppresses the gonadal DTC migration defect of *mig-17(tk174)* null mutants. The *rpl-20* mutants grew slower than wild-type animals. The slow-growing mutants, *clk-1* or *clk-2*, also suppressed *mig-17*, albeit partially, suggesting that an additional mechanism acts in the suppression by *rpl-20(tk73)*. We found that *rpl-20(tk73)* requires FBL-1C and NID- 1, the downstream target of MIG-17 in the regulation of DTC migration, to suppress *mig-17* mutants and that the amount of basement membrane accumulation of FBL-1C was upregulated in *rpl-20(tk73)* mutants.

It is surprising that a mutation in a ribosome subunit strongly suppresses the gonadal defect of mutations in *mig-17*, which encodes a matrix metalloprotease that acts in remodeling the basement membrane. The ribosome is a cellular factory comprising multiple ribosomal RNAs and proteins and functions in the translational production of proteins from mRNAs. Although mutations in ribosomal proteins can affect translational efficacy in general, defects in ribosome biogenesis caused by mutations in single ribosomal protein genes result in tissue-specific phenotypic abnormalities. In humans, mutations in nineteen out of eighty-one ribosomal protein genes, not including RPL18a/eL20 thus far, are known to be causative for dominant genetic disorders called ribosomopathies. Ribosomopathies are mainly characterized by erythroid hypoplasia in the bone marrow (Diamond-Blackfan anemia; DBA), but they also associate with a series of phenotypic spectra, including craniofacial, limb, cardiac, and genitourinary malformations, as well as growth retardation and increased risks of cancers. DBA is caused by inhibition of the hematopoietic system through nucleolar stress-dependent stabilization of p53, which leads to cell cycle arrest and apoptosis (TENG *et al*. 2013; KANG *et al*. 2021). DBA patients with mutations in RPL9 exhibit a reduction of the 60S subunit coupled with a reduction of the 80S ribosomes, which is reminiscent of our observation in *rpl-20(tk73)* (LEZZERINI et al. 2020).

The mechanisms of abnormality in organogenesis have been better characterized in mice and *Drosophila*. Belly spot and tail (*Bst*) is a semi-dominant, homozygous lethal mutation in RPL24, and *Bst/+* mice have decreased pigmentation, a kinked tail, a reduced number of retinal ganglion cells, and an extra preaxial digit (OLIVER *et al*. 2004). Tail short (*Ts*) is a dominant, homozygous lethal mutation in RPL38, and *Ts/+* animals exhibit a short and kinked tail that is associated with skeletal patterning defects.

Vertebrae extending along the anterior-posterior body axis exhibit homeotic transformations. Interestingly, a specific subset of Hox genes (*Hoxa4*, *a9*, *a11*, *b3*, *c8*, and *d10*) specifying the vertebral patterning are markedly decreased in polysome association in *Ts/+* embryos, suggesting that RPL38 has a specialized function in translational control of these mRNAs (KONDRASHOV *et al*. 2011). A *Drosophila* model carrying a mutation in the 40S subunit *RPS23^R67K/+^*, corresponding to one of the human ribosomopathies, exhibits scutellar bristles and delayed larval development. Molecular analysis of *RPS23^R67K/+^*animals revealed that the stoichiometric imbalance of protein components of the 40S subunit triggers proteotoxic stress, which decreases ribosomal protein production, slows down protein synthesis, and stimulates autophagy through inhibition of the TOR pathway (RECASENS-ALVAREZ *et al*. 2021).

Although the reason for slow growth in *rpl-20(tk73)* mutants is unknown, growth retardation is a common feature of ribosomopathies. In *mig-17* mutants, the DTCs often detach from the body wall, their natural substratum, and instead attach to the intestine or gonad and migrate over these tissues (NISHIWAKI *et al*. 2000). This suggests that the activity of MIG-17 is important for the adhesion of DTCs to the body wall, which may lead to appropriate responses to guidance molecules such as UNC-6/netrin and Wnt (LEVY-STRUMPF AND CULOTTI 2014; LEVY-STRUMPF *et al*. 2015). When *mig-17(k174)* mutants are cultured at 16°C, the growth rate is delayed by about 1.5 times (BYERLY *et al*. 1976), and the DTC phenotype is weakened (DTC migration defect: 37% anterior and 72% posterior at 20°C, 18% anterior and 53% posterior at 16°C). Since *k174* is a null mutation, the *mig-17* gene function cannot be restored even at low temperatures. As the penetrance of *mig-17(k174)* mutants with an abnormal DTC migration phenotype is around 70 to 80%, it is considered that a second pathway regulates DTC migration, besides the MIG-17 pathway. One possibility is that at 16°C, the duration for adhesion to the body wall and response to guidance molecules is prolonged, which may allow more time for the second pathway to function in the absence of MIG-17. It might be possible that a similar situation occurs in the *rpl-20(tk73)* mutants even at 20°C.

Why was the amount of FBL-1C localized to the basement membrane increased, although the expressed protein level of FBL-1C was not increased in *rpl-20(tk73)*? It is possible that the basement membrane has a higher affinity for FBL-1C in the *rpl- 20(tk73)* background. Since FBL-1C is secreted from the intestine into the body cavity and then localized to the basement membrane, the expression level of receptor molecules present in the basement membrane may be increased, or molecules that decrease the affinity of FBL-1C may be suppressed. Overexpression of the *rpl-20(tk73)* mutant gene in the intestine most strongly suppressed DTC migration abnormalities in *mig-17* mutants, but overexpression in muscle, pharynx, and DTCs also had a weak suppressive effect. These results suggest that such FBL-1C affinity-regulating molecules are secreted. The *rpl-20(tk73)* mutant protein may promote the translation of the receptor molecule or suppress the translation of the affinity-lowering molecule.

Interestingly, FBL-1C can rescue *fbl-1* null mutants when expressed in the intestine but not when expressed outside the intestine (MURIEL *et al*. 2005). This fact suggests the possibility that FBL-1C is secreted from the intestine as a complex with some unknown molecule important for FBL-1C function. It might be possible that *rpl-20(tk73)* influences the translation of an unknown protein that forms a complex with FBL-1C, which functions in the modulation of FBL-1C’s affinity to the basement membrane.

As reported in mammals and *Drosophila*, it might be possible that stress responses involving p53 stabilization or TOR pathway downregulation are triggered by the ribosomal mutation *rpl-20(tk73)*. Further molecular analysis will be needed to understand the precise mechanism downstream of *rpl-20(tk73)*.

## Data Availability

Strains and plasmids are available upon request. The authors affirm that all data necessary for confirming the conclusions of the article are present within the article and figures.

## Acknowledgments

We thank Chizu Yoshikata, Nami Okahashi, Noriko Nakagawa and Asami Sumitani for technical assistance. Some nematode strains used in this work were provided by the Caenorhabditis Genetics Center, which is funded by the National Institutes of Health National Center for Research Resources and from Shohei Mitani through the National Bioresource Project for the nematode.

## Funding

This work was supported by a Grant-in-Aid for Challenging Research by Ministry of Education, Culture, Sports, Science and Technology to KN (26650086).

## Supporting Information

**Figure S1.**
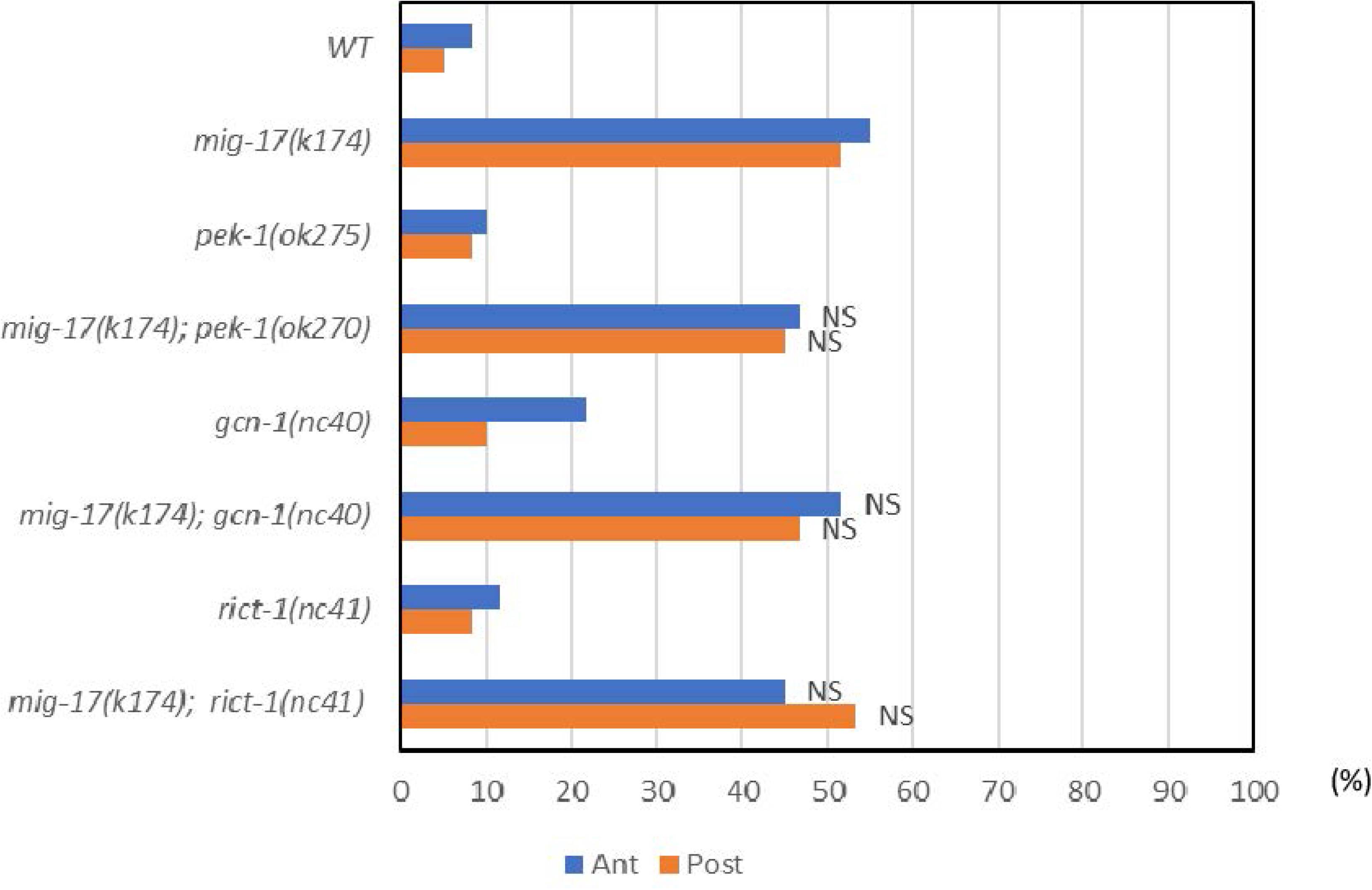
Quantitative analysis of gonadal defects for *gcn-1; mig-17*, *pek-1; mig-17* and *rict-1; mig-17*. *gcn-1*, *pek-1* and *rict-1* failed to suppress *mig-17. mig-17(k174)* was marked with *unc-42(e270),* which weakens *mig-17* phenotype. N=60 for each strain. Significance for Fisher’s exact test against *mig-17(k174)* are indicated: NS, not significant.

**Figure S2.**
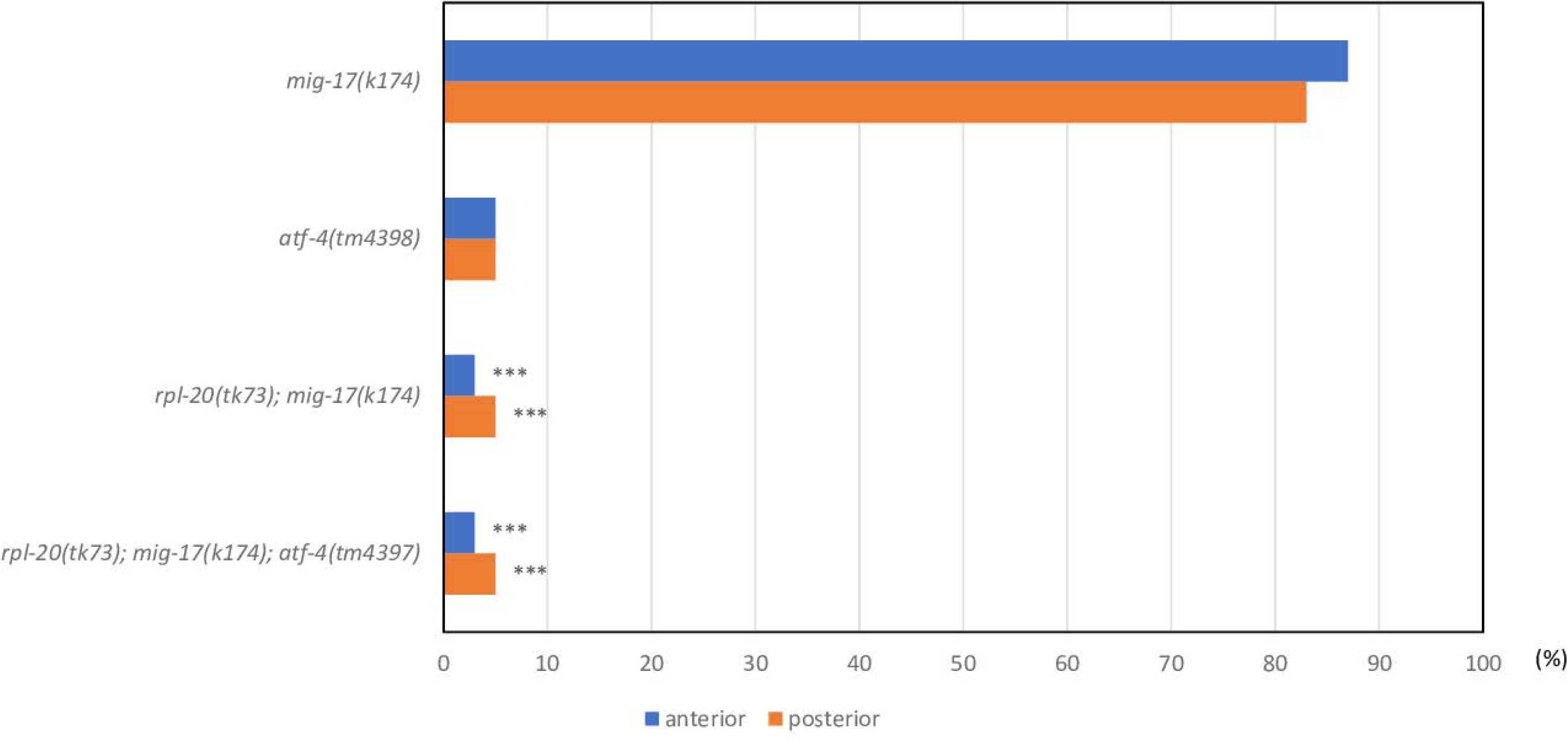
Quantitative analysis of gonadal defects for mig-17(k174), atf-4(tm4398), rpl-20((tk73); mig- 17(k174) and rpl-20(tk73); mig-17(k174); atf-4(tm4397). atf-4(tm4397) does not affect the suppressor activity of rpl-20(tk73). N=60 for each strain. P-values for Fisher’s exact test against mig-17(k174) for mig-17(k174) carrying strains: ***, P < 0.001.

## Literature Cited

Ahmed, S., A. Alpi, M. O. Hengartner and A. Gartner, 2001 C. elegans RAD-5/CLK-2 defines a new DNA damage checkpoint protein. Curr Biol 11: 1934–1944.

Apte, S. S., 2009 A disintegrin-like and metalloprotease (reprolysin-type) with thrombospondin type 1 motif (ADAMTS) superfamily: functions and mechanisms. J Biol Chem 284: 31493–31497.

Ban, N., R. Beckmann, J. H. Cate, J. D. Dinman, F. Dragon et al., 2014 A new system for naming ribosomal proteins. Curr Opin Struct Biol 24: 165–169.

Benard, C., B. McCright, Y. Zhang, S. Felkai, B. Lakowski et al., 2001 The C. elegans maternal-effect gene clk-2 is essential for embryonic development, encodes a protein homologous to yeast Tel2p and affects telomere length. Development 128: 4045–4055.

Brenner, S., 1974 The genetics of Caenorhabditis elegans. Genetics 77: 71–94.

Byerly, L., S. Scherer and R. L. Russell, 1976 The life cycle of the nematode Caenorhabditis elegans. II. A simplified method for mutant characterization. Dev Biol 51: 34–48.

Enomoto, H., C. M. Nelson, R. P. Somerville, K. Mielke, L. J. Dixon et al., 2010 Cooperation of two ADAMTS metalloproteases in closure of the mouse palate identifies a requirement for versican proteolysis in regulating palatal mesenchyme proliferation. Development 137: 4029–4038.

Fujiwara, T., K. Ito and Y. Nakamura, 2001 Functional mapping of ribosome-contact sites in the ribosome recycling factor: a structural view from a tRNA mimic. RNA 7: 64–70.

Fukushige, T., M. G. Hawkins and J. D. McGhee, 1998 The GATA-factor elt-2 is essential for formation of the Caenorhabditis elegans intestine. Dev Biol 198: 286–302.

Guo, Y., C. Tocchini and R. Ciosk, 2021 CLK-2/TEL2 is a conserved component of the nonsense-mediated mRNA decay pathway. PLoS One 16: e0244505.

Hesselson, D., C. Newman, K. W. Kim and J. Kimble, 2004 GON-1 and fibulin have antagonistic roles in control of organ shape. Curr Biol 14: 2005–2010.

Ihara, S., and K. Nishiwaki, 2007 Prodomain-dependent tissue targeting of an ADAMTS protease controls cell migration in Caenorhabditis elegans. EMBO J 26: 2607–2620.

Imanishi, A., Y. Aoki, M. Kakehi, S. Mori, T. Takano et al., 2020 Genetic interactions among ADAMTS metalloproteases and basement membrane molecules in cell migration in Caenorhabditis elegans. PLoS One 15: e0240571.

Kang, J., N. Brajanovski, K. T. Chan, J. Xuan, R. B. Pearson et al., 2021 Ribosomal proteins and human diseases: molecular mechanisms and targeted therapy. Signal Transduct Target Ther 6: 323.

Keeley, D. P., E. Hastie, R. Jayadev, L. C. Kelley, Q. Chi et al., 2020 Comprehensive Endogenous Tagging of Basement Membrane Components Reveals Dynamic Movement within the Matrix Scaffolding. Dev Cell 54: 60–74 e67.

Kim, H. S., Y. Kitano, M. Mori, T. Takano, T. E. Harbaugh et al., 2014 The novel secreted factor MIG-18 acts with MIG-17/ADAMTS to control cell migration in Caenorhabditis elegans. Genetics 196: 471–479.

Kondrashov, N., A. Pusic, C. R. Stumpf, K. Shimizu, A. C. Hsieh et al., 2011 Ribosome-mediated specificity in Hox mRNA translation and vertebrate tissue patterning. Cell 145: 383–397.

Kubota, Y., R. Kuroki and K. Nishiwaki, 2004 A fibulin-1 homolog interacts with an ADAM protease that controls cell migration in C. elegans. Curr Biol 14: 2011–2018.

Kubota, Y., K. Nagata, A. Sugimoto and K. Nishiwaki, 2012 Tissue architecture in the Caenorhabditis elegans gonad depends on interactions among fibulin-1, type IV collagen and the ADAMTS extracellular protease. Genetics 190: 1379–1388.

Kubota, Y., K. Ohkura, K. K. Tamai, K. Nagata and K. Nishiwaki, 2008 MIG-17/ADAMTS controls cell migration by recruiting nidogen to the basement membrane in C. elegans. Proc Natl Acad Sci U S A 105: 20804–20809.

Lakowski, B., and S. Hekimi, 1996 Determination of life-span in Caenorhabditis elegans by four clock genes. Science 272: 1010–1013.

Levy-Strumpf, N., and J. G. Culotti, 2014 Netrins and Wnts function redundantly to regulate antero-posterior and dorso-ventral guidance in C. elegans. PLoS Genet 10: e1004381.

Levy-Strumpf, N., M. Krizus, H. Zheng, L. Brown and J. G. Culotti, 2015 The Wnt Frizzled Receptor MOM-5 Regulates the UNC-5 Netrin Receptor through Small GTPase- Dependent Signaling to Determine the Polarity of Migrating Cells. PLoS Genet 11: e1005446.

Lezzerini, M., M. Penzo, M. F. O’Donohue, C. Marques Dos Santos Vieira, M. Saby, et al., 2020 Ribosomal protein gene RPL9 variants can differentially impair ribosome function and cellular metabolism. Nucleic Acids Res 48: 770–787.

Maduro, M., and D. Pilgrim, 1995 Identification and cloning of unc-119, a gene expressed in the Caenorhabditis elegans nervous system. Genetics 141: 977–988.

McCulloch, D. R., C. M. Nelson, L. J. Dixon, D. L. Silver, J. D. Wylie et al., 2009 ADAMTS metalloproteases generate active versican fragments that regulate interdigital web regression. Dev Cell 17: 687–698.

Mittaz, L., D. L. Russell, T. Wilson, M. Brasted, J. Tkalcevic et al., 2004 Adamts-1 is essential for the development and function of the urogenital system. Biol Reprod 70: 1096–1105.

Muriel, J. M., C. Dong, H. Hutter and B. E. Vogel, 2005 Fibulin-1C and Fibulin-1D splice variants have distinct functions and assemble in a hemicentin-dependent manner. Development 132: 4223–4234.

Nishiwaki, K., 1999 Mutations affecting symmetrical migration of distal tip cells in Caenorhabditis elegans. Genetics 152: 985–997.

Nishiwaki, K., N. Hisamoto and K. Matsumoto, 2000 A metalloprotease disintegrin that controls cell migration in Caenorhabditis elegans. Science 288: 2205–2208.

Nukazuka, A., H. Fujisawa, T. Inada, Y. Oda and S. Takagi, 2008 Semaphorin controls epidermal morphogenesis by stimulating mRNA translation via eIF2alpha in Caenorhabditis elegans. Genes Dev 22: 1025–1036.

Nukazuka, A., S. Tamaki, K. Matsumoto, Y. Oda, H. Fujisawa et al., 2011 A shift of the TOR adaptor from Rictor towards Raptor by semaphorin in C. elegans. Nat Commun 2: 484.

Okkema, P. G., S. W. Harrison, V. Plunger, A. Aryana and A. Fire, 1993 Sequence requirements for myosin gene expression and regulation in Caenorhabditis elegans. Genetics 135: 385–404.

Oliver, E. R., T. L. Saunders, S. A. Tarle and T. Glaser, 2004 Ribosomal protein L24 defect in belly spot and tail (Bst), a mouse Minute. Development 131: 3907–3920.

Recasens-Alvarez, C., C. Alexandre, J. Kirkpatrick, H. Nojima, D. J. Huels et al., 2021 Ribosomopathy-associated mutations cause proteotoxic stress that is alleviated by TOR inhibition. Nat Cell Biol 23: 127–135.

Robert, V. J., T. Sijen, J. van Wolfswinkel and R. H. Plasterk, 2005 Chromatin and RNAi factors protect the C. elegans germline against repetitive sequences. Genes Dev 19: 782–787.

Russell, D. L., K. M. Doyle, S. A. Ochsner, J. D. Sandy and J. S. Richards, 2003 Processing and localization of ADAMTS-1 and proteolytic cleavage of versican during cumulus matrix expansion and ovulation. J Biol Chem 278: 42330–42339.

Saijou, E., T. Fujiwara, T. Suzaki, K. Inoue and H. Sakamoto, 2004 RBD-1, a nucleolar RNA-binding protein, is essential for Caenorhabditis elegans early development through 18S ribosomal RNA processing. Nucleic Acids Res 32: 1028–1036.

Shindo, T., H. Kurihara, K. Kuno, H. Yokoyama, T. Wada et al., 2000 ADAMTS-1: a metalloproteinase-disintegrin essential for normal growth, fertility, and organ morphology and function. J Clin Invest 105: 1345–1352.

Simsek, D., and M. Barna, 2017 An emerging role for the ribosome as a nexus for post- translational modifications. Curr Opin Cell Biol 45: 92–101.

Suzuki, N., H. Toyoda, M. Sano and K. Nishiwaki, 2006 Chondroitin acts in the guidance of gonadal distal tip cells in C. elegans. Dev Biol 300: 635–646.

Tamai, K. K., and K. Nishiwaki, 2007 bHLH transcription factors regulate organ morphogenesis via activation of an ADAMTS protease in C. elegans. Dev Biol 308: 562–571.

Teng, T., G. Thomas and C. A. Mercer, 2013 Growth control and ribosomopathies. Curr Opin Genet Dev 23: 63–71.

Wicks, S. R., R. T. Yeh, W. R. Gish, R. H. Waterston and R. H. Plasterk, 2001 Rapid gene mapping in Caenorhabditis elegans using a high density polymorphism map. Nat Genet 28: 160–164.

Wong, A., P. Boutis and S. Hekimi, 1995 Mutations in the clk-1 gene of Caenorhabditis elegans affect developmental and behavioral timing. Genetics 139: 1247–1259.

